# Insights into tick-pathogen interactions – a single cell RNA sequencing approach of transcriptional changes during ehrlichial infection

**DOI:** 10.64898/2026.03.19.712879

**Authors:** Abdulsalam Adegoke, Joseph Aspinwall, Colton McNinch, Margaret C. W. Ho, Adam X. Miranda, Forrest Hoyt, Vinod Nair, Justin Lack, Tais Berelli Saito

**Affiliations:** Laboratory of Malaria and Vector Research, Division of Intramural Research, National Institute of Allergy and Infectious Diseases, National Institutes of Health, Rockville, Maryland 20852, USA; Bioinformatics and Computational Biosciences, Office of Cyber Infrastructure and Computational Biology, Division of Intramural Research, National Institute of Allergy and Infectious Diseases, National Institutes of Health, Bethesda, MD 20892; Research Technologies Branch, Division of Intramural Research, National Institute of Allergy and Infectious Diseases, National Institutes of Health, Bethesda, MD 20892; Current at the Laboratory of virology, Division of Intramural Research, National Institute of Allergy and Infectious Diseases, National Institutes of Health, Hamilton, MT 59840

**Keywords:** Tick, *Ixodes scapularis*, ISE6, *Ehrlichia muris eauclairensis*, single cell, RNA-Sequencing, vector-pathogen interaction

## Abstract

Tick-borne diseases represent a significant threat to human and animal health worldwide. In the United States, the blacklegged tick, *Ixodes scapularis* (*I. scapularis)*, serves as a competent vector for several bacterial pathogens, including *Ehrlichia muris eauclairensis* (EME). The *I. scapularis* embryonic cell line (ISE6) is a valuable tool for propagating tick-borne pathogens and studying tick-pathogen interactions. In this study, we examined the cellular complexity of ISE6 cells and their response to EME infection. Single-cell RNA sequencing revealed 15 distinct cell clusters present. Although ISE6 cells are heterogeneous, they do not display transcriptional similarity to any known tick tissues. Notably, this lack of similarity did not influence their susceptibility to EME infection. Our results demonstrated that EME infection induces time-dependent transcriptional changes in ISE6 cells: early infection is characterized by upregulation of genes associated with stress adaptation, mitochondrial function, and metabolic pathways, whereas late infection leads to broad downregulation of genes involved in the cell cycle, DNA replication, and cytoskeletal organization. These findings enhance our understanding of ehrlichial interactions with ISE6 cells and reinforce the utility of this cell line as a resource for isolating and propagating arthropod endosymbionts and tick-borne pathogens.

**IMPORTANCE:** This study provides a single-cell resolution framework for interpreting tick cell line biology during infection with a medically relevant ehrlichial pathogen. Using scRNA-seq, we show that the *I. scapularis* embryonic-derived ISE6 cell line comprises multiple transcriptionally distinct cell states, yet these states do not map cleanly onto canonical tick tissue signatures, even when compared against a curated reference tissue atlas. Despite this heterogeneity, EME broadly infects ISE6 cell population, indicating that susceptibility is not restricted to a specific cell type. We further define a time-dependent arthropod vector response in which early infection is marked by activation of stress and metabolic adaptation response, followed by late-stage inhibition of key signaling, transcriptional, and proliferative pathways as bacterial burden increases. Together, these findings strengthen the biological interpretation of ISE6 as an in vitro model for tick–pathogen interactions and provide a resource for future mechanistic studies of ehrlichial persistence, replication, and vector competence.

## INTRODUCTION

Vector-borne diseases are expanding in geographic range and incidence, driven in part by ecological disruption and environmental changes (Paddock et al., 2016). In the United States, *Ixodes scapularis* (*I. scapularis*) serves as a competent vector for multiple bacterial and viral pathogens, including *Ehrlichia muris eauclairensis* (EME), an obligate intracellular bacterium that causes human ehrlichiosis. *Ehrlichia* infection causes an acute febrile illness characterized by hematologic abnormalities and systemic inflammation (Olano et al., 2003; Johnson et al., 2015; Saito et al., 2015a, b). Although not much is known about EME interaction with the vertebrate host (Saito et al., 2015a, b; Ahmed et al., 2022; Migne et al., 2022), even less is known about tick-pathogen interaction (Aspinwall et al., 2025).

Understanding vector–pathogen interactions is critical because pathogen persistence, replication, and transmission depend on the successful colonization of specific tick tissues. EME has been detected in multiple developmental stages of *I. scapularis*, indicating that it persists across the tick life cycle (Lynn et al., 2019). During blood feeding, replication and migration within the tick facilitate transmission to vertebrate hosts and contribute to enzootic maintenance in endemic regions (Lynn et al., 2015; Saito et al., 2015b). However, the molecular mechanisms that govern EME replication and adaptation within tick cells remain poorly defined.

Tick-derived cell lines serve as experimental systems for studying tick–pathogen interactions (Bell-Sakyi et al., 2007; Sharma et al., 2022). Among these, the embryonic-derived ISE6 cell line from *I. scapularis* has been widely used to investigate intracellular pathogens (Munderloh et al., 1994; Villar et al., 2015; Alberdi et al., 2016). ISE6 cells exhibit hemocyte-like morphology and display epithelial and neuronal features (Munderloh et al., 1994; Villar et al., 2015; Alberdi et al., 2016). Molecular profiling studies have also suggested similarities to multiple tick tissues (Oliver et al., 2015; Alberdi et al., 2016; Cabezas-Cruz et al., 2017; Mateos-Hernández et al., 2021).

Although several studies have reported that ISE6 cells reflect specific tissue behaviors, no study has comprehensively compared their transcriptional signatures with those reported for different tissues. In contrast, mosquito cell lines have been more extensively characterized and refined (Barletta et al., 2012; Vasconcellos et al., 2022). These studies highlight the need for comparable analyses in ticks to better define the biological relevance of ISE6 cells and clarify their relationship with EME.

In the present study, we used single-cell RNA sequencing to examine transcriptional heterogeneity in ISE6 cells and their responses to EME infection over time. We evaluated tissue-associated transcriptional signatures and responses to EME infection during early and late stages. These findings clarify the biological properties of ISE6 cells and provide insight into EME–vector interactions.

## MATERIALS AND METHIDS

### Cell culture

ISE6 cells were obtained from Dr. Ulrike Munderloh’s laboratory at the University of Minnesota (St. Paul, MN). Cells were maintained in L15C300 medium as previously described (Munderloh et al., 1994; Munderloh et al. 1999). Cultures were kept at 32°C without CO_2_ and passaged every 7-10 days. For experiments, both uninfected control cells and EME-infected cells were cultured under identical conditions in the same “modified L15C300 medium”, supplemented with 0.25% NaHCO_3_ and 25 mM HEPES, at pH 7,0-7,5, as previously described (Munderloh et al., 1999). Cell smears stained with a modified Giemsa method (HEMA 3, Fisher Scientific), following the manufacturer’s instructions, were used to visualize cellular morphology and infection.

### Bacteria

*Ehrlichia muris* subspecies *eauclairensis* infected ISE6 cells were obtained from Dr. Ulrike Munderloh’s laboratory at the University of Minnesota (St. Paul, MN). The EME infected cells were cultivated with the “modified media” (see details at cell culture section) and maintained in 34°C with 5% CO_2_. The level of ehrlichial infection was qualitatively monitored in smears of the cell monolayer stained with the modified Giemsa dye method (HEMA 3, Fisher Scientific, Waltham, MA).

### Viability and EME quantification in ISE6 cells

After inoculation of ISE6 cells with EME, we collected samples in triplicate at 6, 24, 48, 72, 96, 120 hours post infection (p.i.) to evaluate the bacterial growth in tick cells. The cells were harvested and simultaneously prepared both for flow cytometry to assess the viability of the ISE6 cells and to determine the proportion of infected cells by measuring the expression of ehrlichial surface protein, as well as detection of ehrlichial *dsb* (disulfide oxidoreductase) gene by real-time PCR. To assess both cell viability and the proportion of viable cells infected with EME, samples were stained for dead cells using the live/dead fixable blue kit (Invitrogen, Waltham, MA) following manufacturer’s protocol. The cells were further permeabilized using fixation/permeabilization reagents (BD biosciences, Franklin Lakes, NJ) for 20 minutes at 4°C and incubated with 1:4 dilution of Ec56.5 monoclonal mouse antibody (DSHB, Iowa City, IA) against Ehrlichial OMP-19 surface protein (Li et al., 2001). After incubation overnight at 4°C, the samples were stained with 1:50 dilution of goat anti-mouse FITC-conjugated secondary antibody for one hour at 4°C. The samples were analyzed by flow cytometry to obtain the count of live cells and geometric mean fluorescent intensity (MFI) for ehrlichial OMP-19.

Similarly, DNA was isolated from the other half of the isolated cells following manufacturer’s protocol of the DNeasy tissue and blood isolation kit (QIAGEN, Valencia, CA). DNA was quantified and used for quantitative real-time PCR to determine the amplification of the *E. muris dsb* gene, using primers and probe described previously (Stevenson et al., 2006; Saito et al., 2015 a, b). Bacterial load was determined using qPCR targeting the dsb gene with a plasmid-generated standard curve included on each plate. Copy numbers were normalized to total DNA concentration measured by PicoGreen fluorescence assay.

### Transmission (TEM) and Scanning Electron Microscopy (SEM)

Samples for electron microscopy (EM) were processed as described previously with some modifications (Kennedy et al., 2007; Malachowa et al., 2024). Briefly, the cells for EM were fixed with 2% paraformaldehyde and 2.5% glutaraldehyde in 0.1M sodium cacodylate buffer. For TEM samples, the cells were washed in buffer, post fixed with reduced osmium tetroxide, treated with tannic acid, *en bloc* stained with uranyl acetate replacement stain, dehydrated in graded ethanol series in a Pelco Biowave microwave. The cells were embedded in Epon/Araldite resin and imaged using Talos L120 C transmission electron microscope using a 4Kx4K Ceta CMOS camera (Thermo Fisher Scientific, Hillsboro, OR). For SEM, the fixed cells were adsorbed on a silicon chip, washed with 0.1M sodium cacodylate buffer, post fixed with reduced osmium tetroxide, dehydrated in graded ethanol series and dried using a critical point dryer. The cells were coated with iridium and were imaged on a Hitachi SU 8000 (Hitachi Hi-Tech in the USA, Schaumburg, IL) scanning electron microscope.

Cell pellets of uninfected (either treated with normal and “modified media”) and EME-infected ISE6 at days two and four p.i. were prepared for transmission (TEM) and scanning electron microscopy (SEM).

### Single-cell RNA sequencing of ISE6 cells

Uninfected and EME–infected ISE6 cultures from the same passage were maintained under similar culture conditions and sampled at 2 and 4 days p.i. Two independent experiments were performed ∼30 days apart. Single-cell suspensions were generated by gentle detachment and filtration (40-µm strainer), followed by centrifugation and resuspension in PBS containing 10% fetal bovine serum. Cell counts and viability were determined using a Countess II FL Automated Cell Counter (Thermo Fisher Scientific, Waltham, MA, USA). Single-cell 3′ gene expression libraries were prepared using the Chromium Next GEM Single Cell 3′ Kit v3.1 (10x Genomics) on a Chromium Controller according to the manufacturer’s protocol. Libraries were sequenced on an Illumina HiSeq4000 using paired-end sequencing (26 bp Read 1, 8 bp i7 index, 98 bp Read 2), targeting ∼5,000 cells per sample. FASTQ files were processed using Cell Ranger v7.2.0 and aligned to the *I. scapularis* reference genome (NCBI RefSeq GCF_016920785.2) to generate feature–barcode matrices. Ambient RNA contamination was estimated using decontX and incorporated into downstream filtering. Dimensionality reduction (PCA/UMAP) and graph-based clustering were performed in Seurat v5.3.0, and cluster stability was assessed across resolutions to select parameters for downstream analyses. Cluster marker genes were identified using Wilcoxon rank-sum testing. Differential expressions between conditions were assessed using a pseudobulk approach with DESeq2 (implemented via Seurat). Gene Ontology (GO) enrichment was performed using topGO (weight algorithm; Fisher’s exact test) with *I. scapularis* annotations from VectorBase (see Supplemental Methods for additional details).

### I. scapularis tissue collection and RNA extraction

Unfed *I. scapularis* females were obtained from the Oklahoma State University Tick Rearing Facility. All ticks used in our experiments were maintained in desiccators at a temperature of 21°C–22°C, with approximately 98% humidity and darkness. Tick dissection and tissue isolation was carried out as previously described (Aspinwall et al., 2025). For the isolation of RNA, tissues were suspended in 20μL of QIAzol (Qiagen) and homogenized in the TissueLyser II (Qiagen) at 30 oscillations/second for 1 minute with a 2.8 mm stainless steel bead. RNA extraction was performed using miRNeasy micro kit (Qiagen) following manufacturer instructions with an on-column DNA digestion.

### Tissue specific gene expression

Tissue-specific genes were identified using the Tau specificity index and further filtered using DESeq2-based enrichment testing (see Supplemental Methods for additional details).

## RESULTS

### Uninfected and EME-infected ISE6 cells display pronounced morphological heterogeneity and variability in nuclear characteristics

Brightfield microscopy revealed variations in ISE6 cell morphology (Fig. 1A). Transmission electron microscopy of uninfected ISE6 cells showed variability in both nuclear number and nuclear shape (Fig. 1B). Some cells exhibited a high nuclear-to-cytoplasmic ratio, with the nucleus occupying nearly the entire cytoplasm, whereas others contained a more centrally positioned nucleus. Nuclear electron density also varies among cells. Cells presented as either multinucleated cells or with irregularly shaped nuclei. Cytoplasmic electron density and content varied among cells but did not correlate with nuclear number or nuclear density (Fig. 1B). In infected cells, transmission electron microscopy (TEM) revealed intracytoplasmic EME-containing morulae in multiple cells (Fig. 1C). These morulae varied in size and number and were observed in cells containing either a single nucleus or multiple nuclei (Fig. 1C). The presence of morulae distorted the cytoplasm, making it difficult to distinguish cellular morphological features (Fig. 1C). We performed scanning electron microscopy (SEM) to further assess the surface morphology of EME-infected cells. SEM analysis confirmed that infected cells also exhibited a range of shapes and sizes (Fig. 1D). Collectively, these results indicate that ISE6 cells comprise morphologically distinct groups regardless of infection status and that EME can infect different cell types.

**Figure 1.**
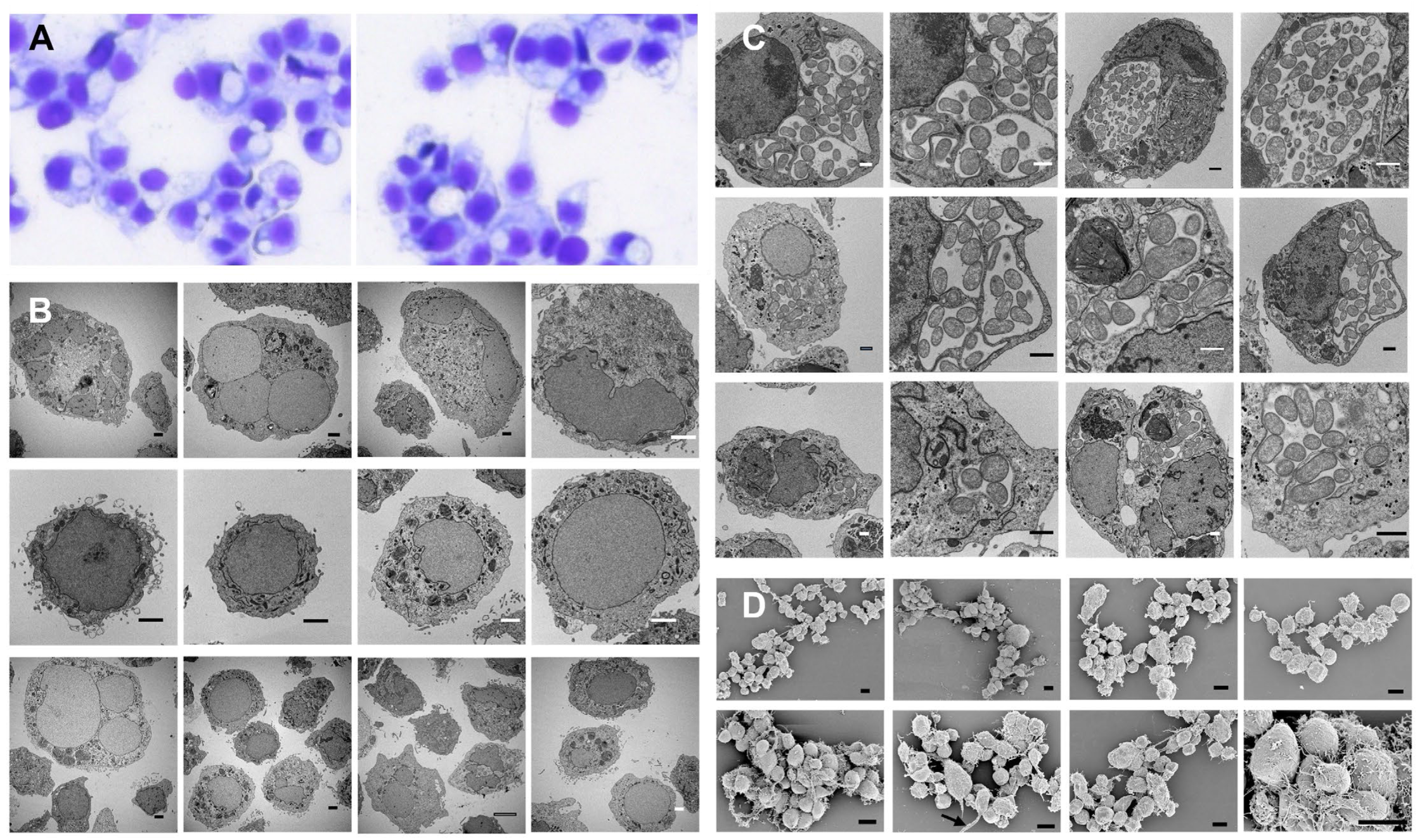
Morphological evaluation of control and EME-infected ISE6 cells. (A) Brightfield photomicrographs of ISE6 cells. (B) Transmission electron microscopy of uninfected ISE6 cells demonstrating differences in cell size, electron density, nuclear size, and nuclear-to-cytoplasmic ratio. (C) Transmission electron microscopy of EME-infected cells showing EME present within single or multiple phagosome-like compartments (morulae) in various cell types. (D) Scanning electron micrographs of ISE6 cells show variation in cell shape and size. Cells also differ in the presence or absence of filopodia-like projections. Scale bars: uninfected TEM = 5 µm; EME-infected TEM = 2 µm; SEM = 50 µm.

### EME replication in ISE6 cells is associated with a progressive decline in host cell viability

To evaluate the impact of EME infection on ISE6 cells, we measured host cell viability and bacterial burden at multiple time points (p.i.). ISE6 cell viability peaked at 48 hours p.i. at 97%, then declined to 82% by 96 hours and fell below 70% at 120 hours, coinciding with the onset of cell monolayer detachment (Fig. 2A; Fig. S1). Quantification of mean fluorescent intensity (MFI) using the ehrlichial OMP-19 antibody showed an initial decrease in bacterial levels at 48 hours, followed by a progressive increase at 96 and 120 hours (p.i.) (Fig. 2A; Fig. S1). This increase in bacterial antigen correlated with a rise in ehrlichial DNA copy numbers, as determined by real-time PCR targeting the *dsb* gene (Fig. 2B). Together, these results demonstrate that EME replicates in the ISE6 cell line over time and that this replication is associated with a decline in host cell viability.

**Figure 2.**
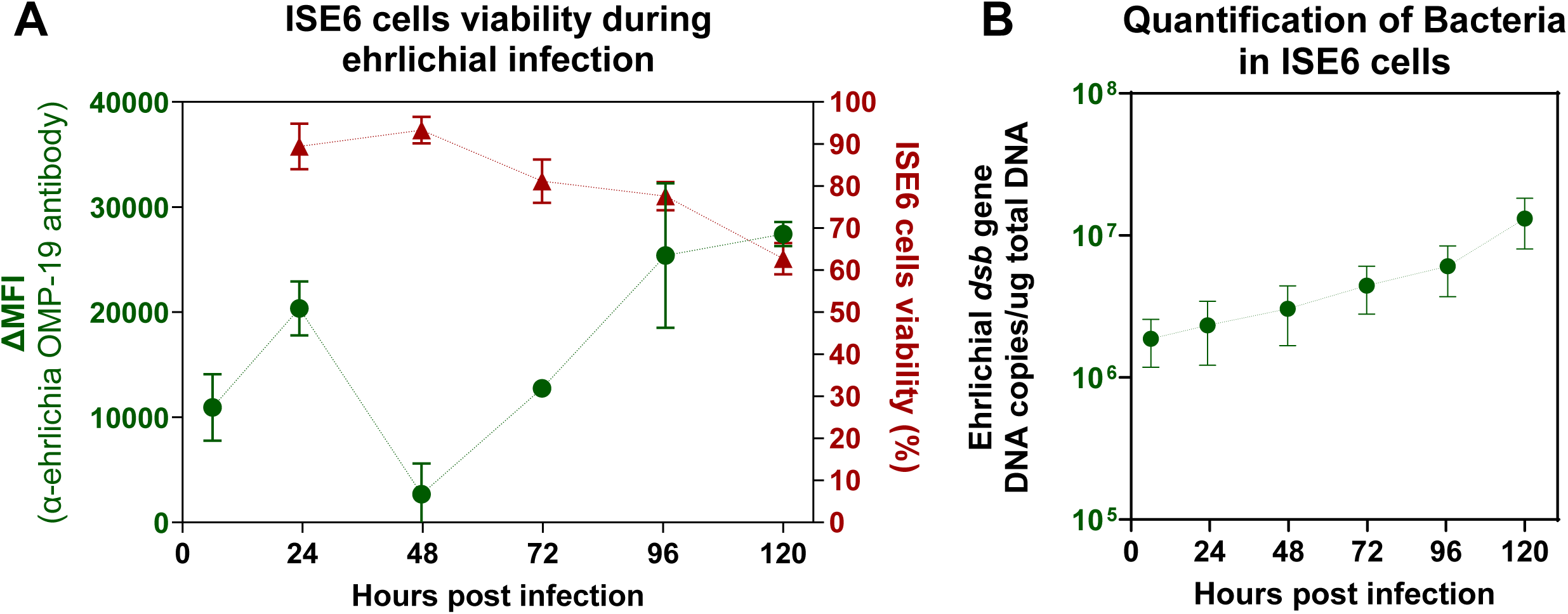
Assessment of ISE6 cells viability in control and EME-infected conditions. (A) The percentage of total viable ISE6 cells (red line) and the mean fluorescence intensity (MFI) of OMP-19 in ISE6 cells (green line), as determined by flow cytometry, throughout the course of EME infection at 0, 24, 28, 72, 96, and 120 hours (p.i.). (B) Quantification of EME *dsb* gene copy number pear µg of DNA in ISE6 cells at 0, 24, 28, 72, 96 and 120 hours (p.i.). Error bars represent standard error of the mean for all samples.

### Defining transcriptional heterogeneity of ISE6 cells using single-cell RNA seq analysis

Single-cell clustering analysis of integrated uninfected and infected samples identified 15 transcriptionally distinct cell clusters at an optimal resolution of 0.2 (Fig. 3A–B; Fig. S2). Putative marker genes were identified for 10 of the 15 clusters (Supplementary Table S1). Cluster distribution analysis showed transcriptional heterogeneity within the ISE6 population (Fig S3 A-B). Several clusters displayed distinct functional signatures. Cluster 6 and Cluster 11 showed enrichment of genes associated with cell cycle regulation and transcriptional control, including transcription factors and signaling molecules. Cluster 13 showed strong enrichment of heat shock proteins, indicating a stress-associated transcriptional profile. Cluster 14 displayed muscle-related signature, including multiple myosin isoforms, troponin I, myocyte-specific enhancer factor 2, and other components of the contractile apparatus. In contrast, several clusters were defined by transport-related or secreted proteins, whereas five clusters lacked detectable marker genes under the applied criteria. Overall, except for muscle-like subpopulation, ISE6 cells did not exhibit strong tissue-specific transcriptional identities, indicating substantial transcriptional heterogeneity without clear differentiation into canonical tissue lineages at single-cell resolution.

**Figure 3.**
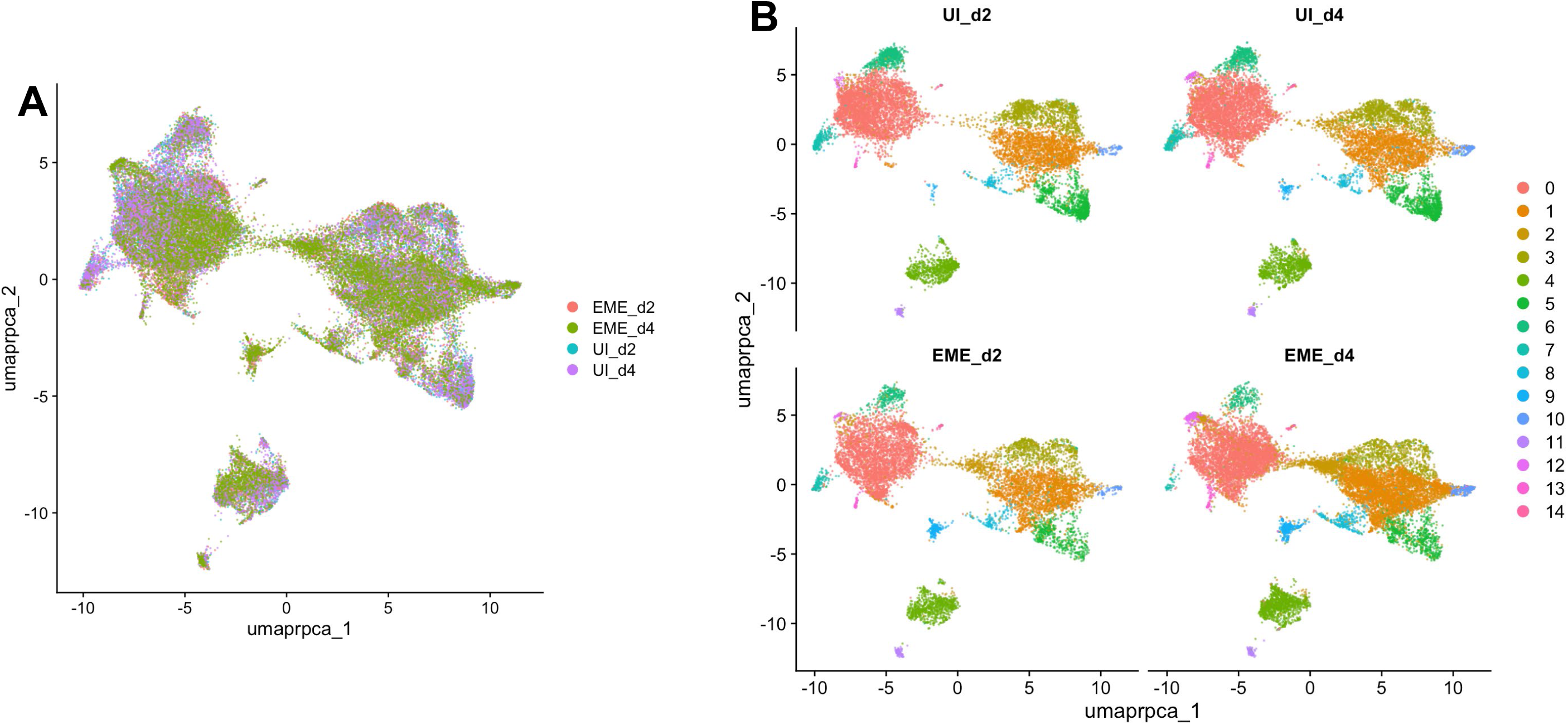
Defining transcriptional heterogeneity of ISE6 cells using single-cell RNA seq analysis. (A and B) Single-cell characterization of uninfected and EME-infected ISE6 cells collected at days-2 and -4 time point post EME infection. (A) Uniform manifold approximation (UMAP) projection demonstrating unsupervised clustering of uninfected and EME-infected ISE6 cells. (B) UMAP projection representing the 15 transcriptional clusters identified across both uninfected and EME-infected ISE6 cells. UMAP was applied on clustering done at a resolution of 0.2.

### EME infection leads to cluster-specific downregulation of functional marker genes in ISE6 cells

Consistent with the increase in EME replication and the decline in host cell viability, we selected two time points for single-cell analysis: day 2 p.i. (EME-d2) (mid-infection, before significant viability loss) and day 4 p.i. (EME-d4), corresponding to peak replication and decreased host cell viability. To determine how EME infection alters cluster-specific markers, we performed differential expression analysis between EME-infected and uninfected ISE6 cells (Fig. 4A–B; Supplementary Table S1). Across multiple clusters, predicted marker genes showed reduced expression in EME-infected cells. Downregulated genes included structural and extracellular matrix–associated components, including hemicentin-2, transport-related genes, transcription factors, stress-associated proteins, and muscle-related genes. Clusters expressing transcriptional regulators (Cluster 11), stress-response genes (Cluster 13), and muscle-associated markers (Cluster 14) showed reduced expression of their respective markers in EME-infected cells compared with uninfected controls. Similarly, clusters defined by secreted or transport-related proteins showed consistent downregulation of their markers in EME-infected cells. Collectively, these results show that EME infection results in the downregulation of functional markers in the ISE6 cells.

**Figure 4.**
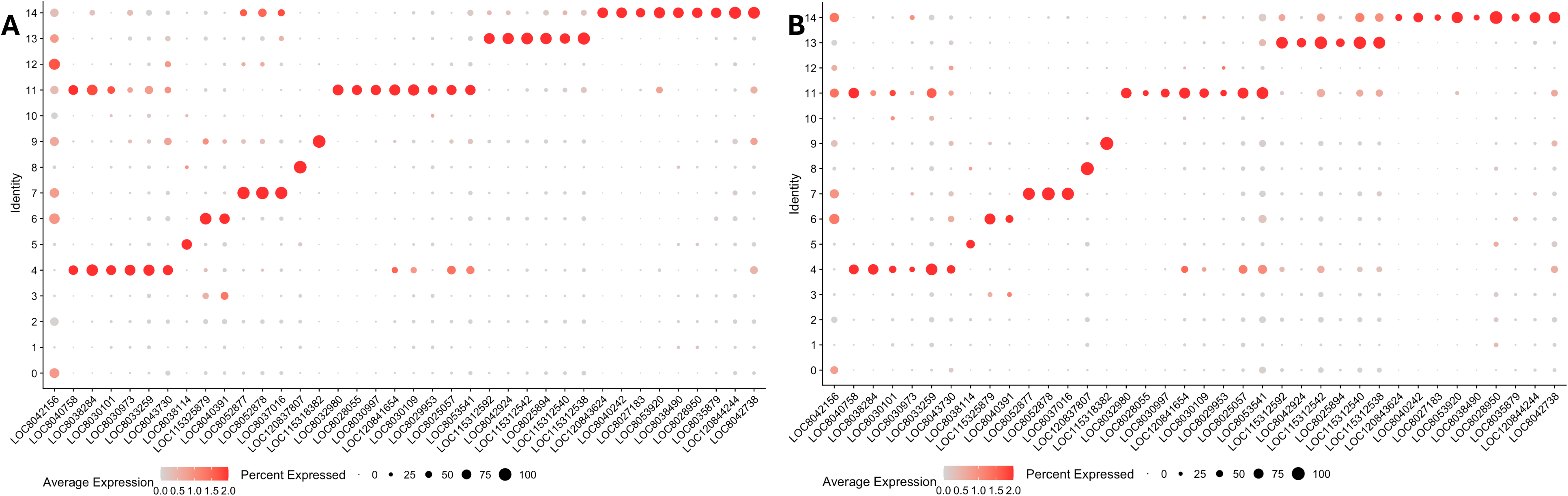
Dot plot of putative transcriptional markers identified across all clusters in uninfected control and EME-infected ISE6 cells. Marker gene expression displayed by dot plot across cell clusters in (A) uninfected and (B) EME-infected ISE6 cells. Dot color shows levels of average expression, while dot size represents the percentage of cells expressing the corresponding genes in each cell cluster.

### ISE6 cells mount an early stress-adaptive transcriptional response to EME infection, which transitions to broad downregulation of metabolic and cell cycle genes as infection progresses

After observing the downregulation of functional ISE6 cell markers in EME-infected cells, we next examined global transcriptional responses across infection stages.

Differential expression analysis revealed EME-driven changes in the ISE6 cell transcriptome, with a greater number of shared differentially expressed genes (DEGs) across clusters in infection-based comparisons than in time-point comparisons within the same condition (Fig. 5A; Supplementary Table S2). Early infection (EME-d2 vs UI-d2) was characterized by upregulated DEGs, whereas later infection (EME-d4 vs UI-d4) showed a shift toward overall downregulation of DEGs, indicating a time-dependent transition from transcriptional activation to suppression (Fig. 5B–C; Supplementary Table S2).

**Figure 5.**
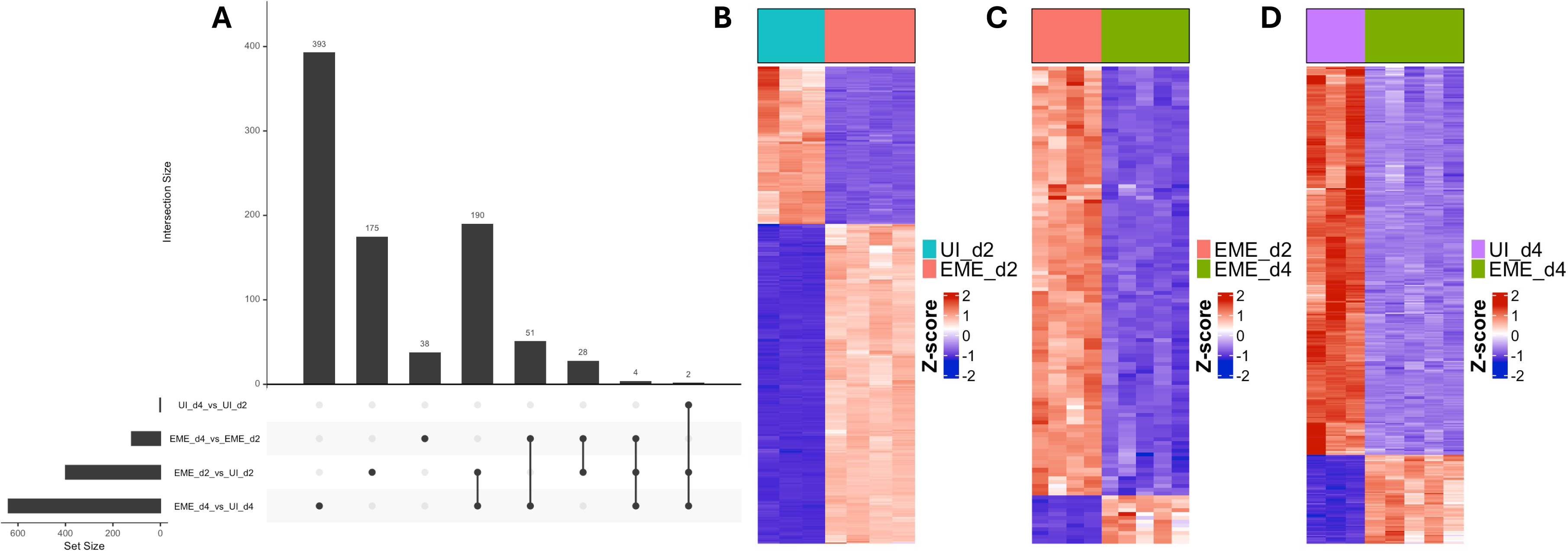
Global transcriptional response of ISE6 cells throughout the course of EME infection. (A) UpSet plot illustrating the overlap of differentially expressed genes (DEGs) among pairwise comparisons, highlighting genes that are either shared between conditions or unique to specific comparisons. Bar heights indicate intersection sizes, while horizontal bars show total set sizes for each comparison. Heatmaps show the normalized expression (Z-score) of all DEGs between (B) EME_d2 and UI_d2, (C) EME_d2 and EME_d4 and (D) EME_d4 and UI_d4. Columns represent experimental groups as indicated by the color bars, rows represent genes, and color intensity reflects relative expression (blue = lower, red = higher).

Gene Ontology (GO) enrichment analysis of EME-d2 relative to day 2 uninfected controls (UI-d2) identified significant enrichment of pathways associated with stress response and cellular adaptation (Fig. 6A; Supplementary Table S3; Supplementary Results). These pathways included unfolded protein binding, cellular response to chemical stimulus, response to oxygen-containing compounds, RNA binding, and mitochondrion organization. Many genes within these categories were upregulated, including regulators of endoplasmic reticulum (ER) stress, redox balance, mitochondrial dynamics, and TOR signaling. In contrast, components of the mismatch repair pathway were significantly downregulated on day 2. Overall, these findings support a model in which early EME infection induces transcriptional programs that promote cellular survival and adaptation to EME-induced stress.

**Figure 6.**
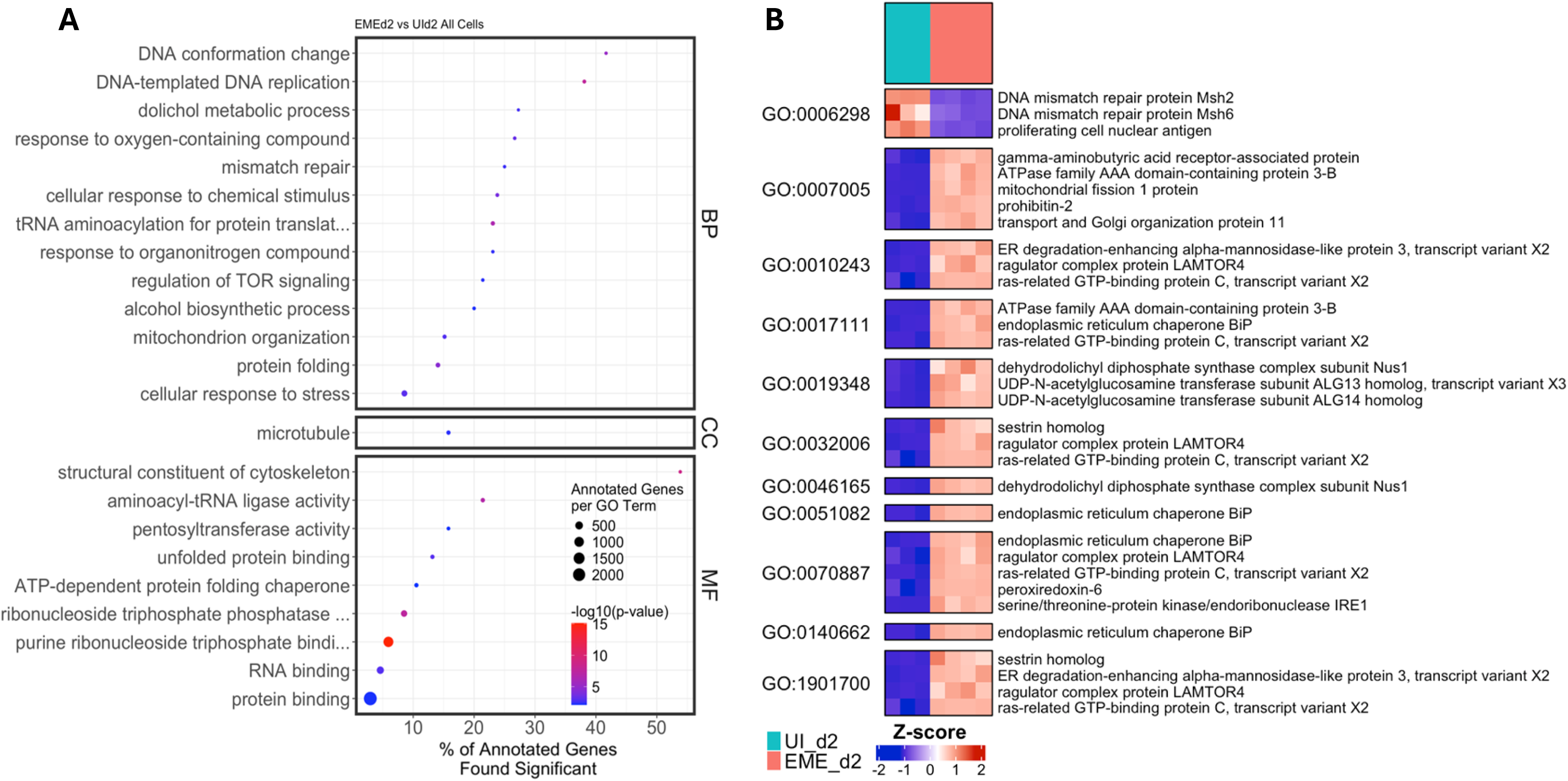
Gene Ontology (GO) and differential expression analyses during early EME infection in ISE6 cells. (A) Dot plots representing GO terms across Biological Process (BP), Cellular Component (CC), and Molecular Function (MF) categories, comparing EME_d2 and UI_d2 ISE6 cells. Dot size indicates the number of annotated genes per GO term, color intensity reflects −log10 (p-value), and the x-axis denotes the percentage of annotated genes that are significantly differentially expressed. (B) Heatmap showing the normalized expression (Z-score) of DEGs within selected enriched GO terms.

Comparison of EME-d 4 and EME-d2 cells revealed enrichment of pathways related to kinase activity, phosphotransferase activity, fatty acid biosynthesis, and mitochondrial outer membrane function (Fig. 7A; Supplementary Table S4; Supplementary Result). These pathways consisted predominantly of downregulated genes at day 4, indicating progressive suppression of signaling cascades, lipid metabolism, and mitochondrial regulatory processes as infection progressed. This shift indicates that the early adaptive response is not sustained but instead transitions to broader attenuation of metabolic and regulatory activity during later stages of infection.

**Figure 7.**
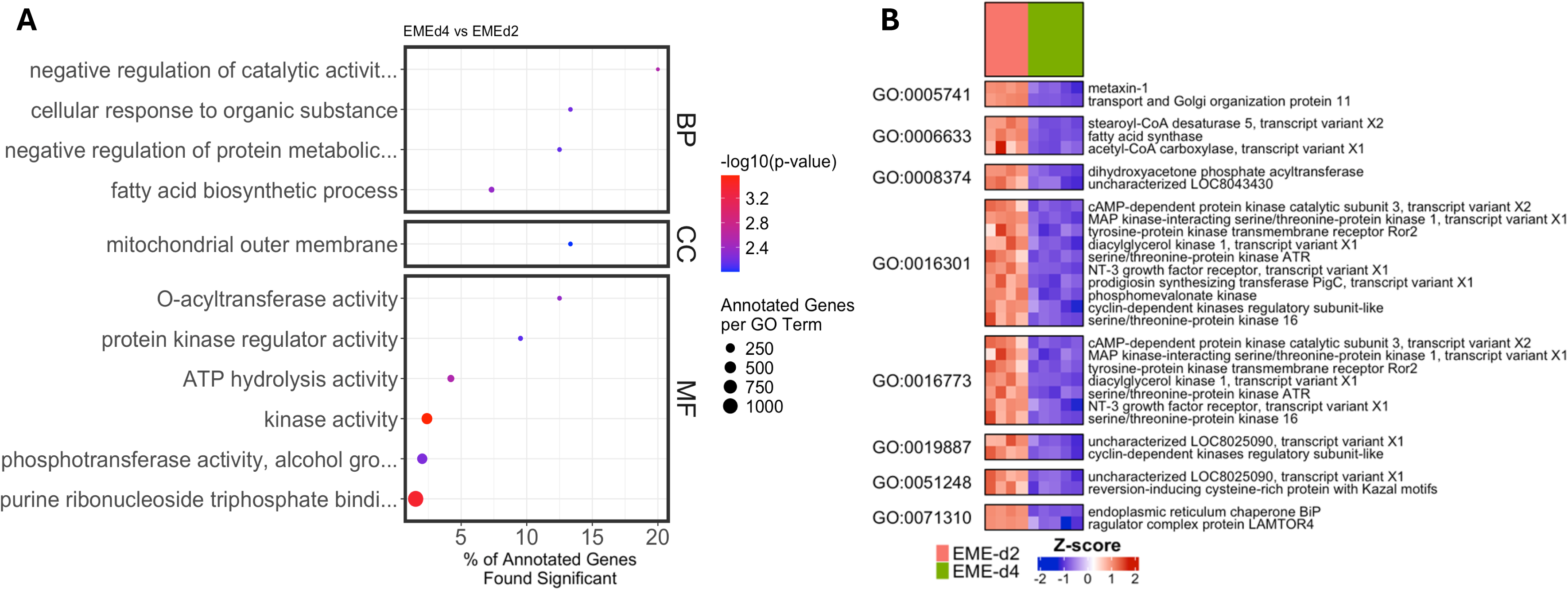
Gene Ontology (GO) and differential expression analyses comparing early and late EME infection in ISE6 cells. (A) Dot plots representing GO terms across Biological Process (BP), Cellular Component (CC), and Molecular Function (MF) categories, comparing EME_d4 and EME_d2 ISE6 cells. Dot size indicates the number of annotated genes per GO term, color intensity reflects −log10 (p-value), and the x-axis denotes the percentage of annotated genes that are significantly differentially expressed. (B) Heatmap showing the normalized expression (Z-score) of DEGs within selected enriched GO terms.

Consistent with this trend, analysis of EME-d4 compared to day 4 uninfected controls (UI-d4) showed significant enrichment of pathways associated with cell division, DNA replication, microtubule organization, and cell cycle checkpoint signaling, suggesting enhanced proliferative and cell cycle activity at this stage of infection (Fig. 8A; Supplementary Table S5; Supplementary Result). These enriched categories were composed predominantly of downregulated genes, including key components of the DNA replication machinery, mitotic checkpoint regulators, DNA repair proteins, and cytoskeletal elements. The suppression of DNA replication factors, MCM complex components, cyclin-dependent kinases, and microtubule-associated proteins indicates inhibition of proliferative and mitotic processes during late infection. Together, these findings suggest that EME infection ultimately drives ISE6 cells toward a state of reduced proliferative capacity and diminished cytoskeletal dynamics. To understand how EME changes gene regulation, we measured transcription factor expression across clusters using composite scores and compared regulatory activity between conditions. (Fig. 9A–C; Supplementary Table S6). Uninfected cells showed higher transcription factor scores compared to infected cells on both day 2 and day 4. This decrease in transcription factor expression further supports the gradual reduction in overall gene activity during infection.

**Figure 8.**
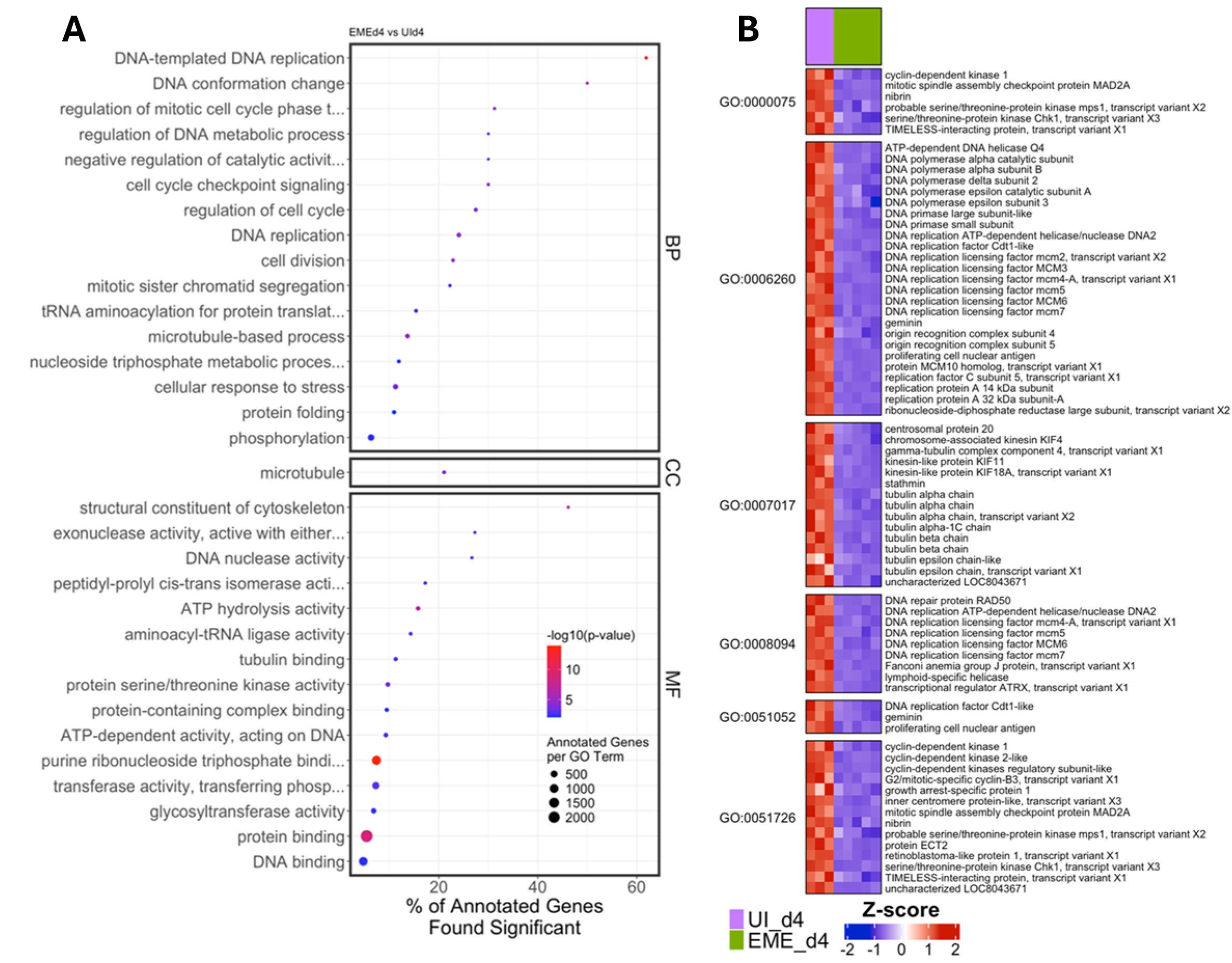
Gene Ontology (GO) and differential expression analyses during late EME infection in ISE6 cells. (A) Dot plots representing GO terms across Biological Process (BP), Cellular Component (CC), and Molecular Function (MF) categories, comparing EME_d4 and UI_d4 ISE6 cells. Dot size indicates the number of annotated genes per GO term, color intensity reflects −log10 (p-value), and the x-axis denotes the percentage of annotated genes that are significantly differentially expressed. (B) Heatmap showing the normalized expression (Z-score) of DEGs within selected enriched GO terms.

**Figure 9.**
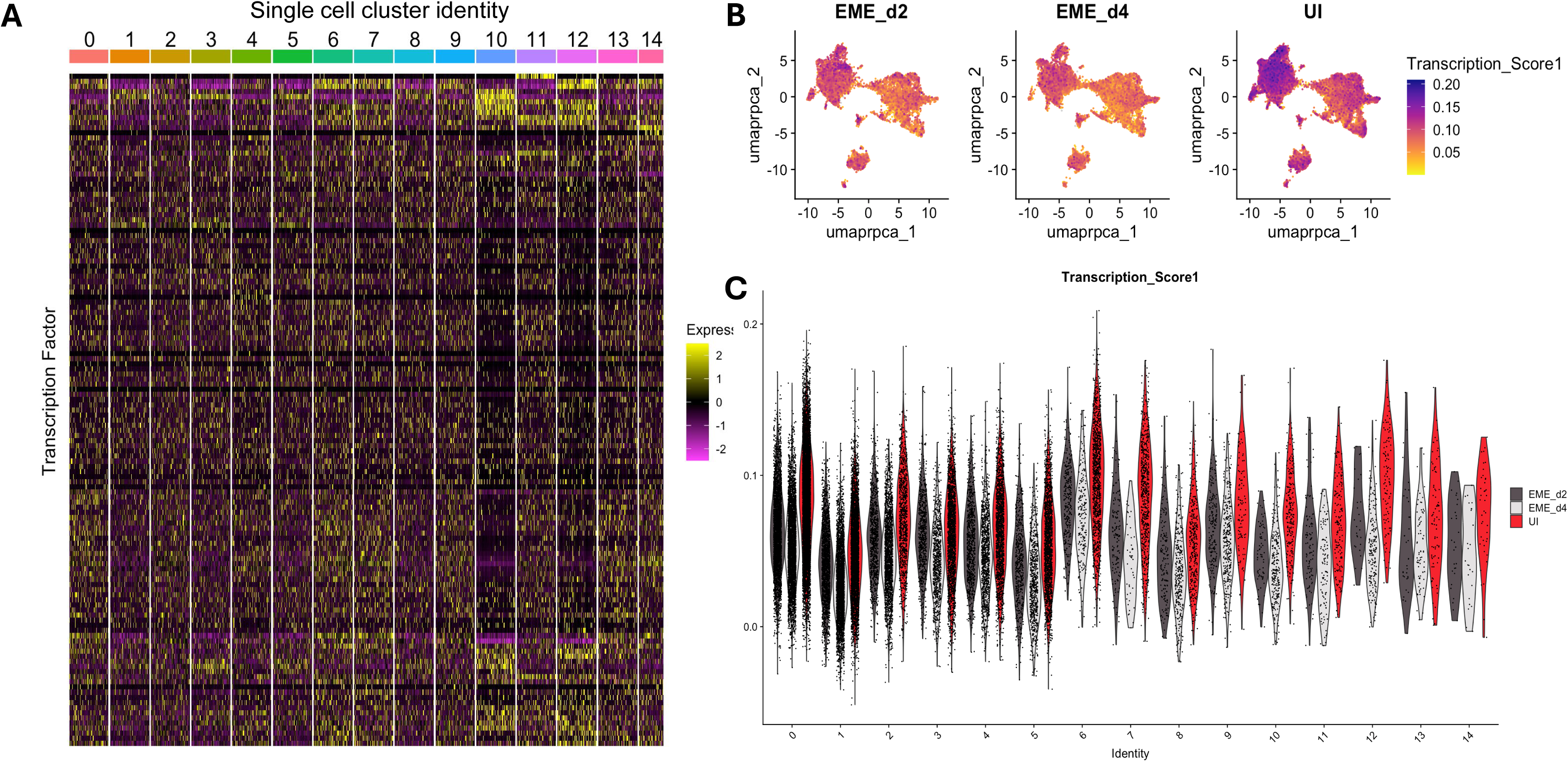
Expression patterns of transcriptional factors in uninfected (UI) and EME-infected (EME) groups. (A) Heatmap showing the normalized expression of transcription factors identified in all the single-cell clusters in the ISE6 cells. (B and C) UMAP projection and violin plot of the transcription score in all single-cell clusters.

### Absence of cell specific and tissue specific signature in ISE6 at single cell resolution

Given the EME-associated transcriptional responses in ISE6 cells, and the fact that several identified markers have defined tissue-specific functions, we next sought to determine if these markers correspond to any specific tissues. We then created a comprehensive tissue transcriptional reference atlas. The atlas included Malpighian tubules, midgut, ovaries, salivary glands, synganglion, trachea, and carcass, while hemocyte expression profiles were taken from published datasets (Rolandelli et al., 2024). For each tissue, we identified genes that were uniquely expressed and used them as tissue-specific markers. These gene sets were then used to map the single-cell ISE6 transcriptome and assign tissue signatures. Although there was transcriptional variation across ISE6 clusters, no cluster showed clear enrichment for a specific tissue signature (Fig. 10A; Supplementary Table S7). Marker genes corresponded to multiple reference tissues, including a leucine-rich repeat protein in hemocytes, sulfotransferase 1E1 in the Malpighian tubules, solute carrier organic anion transporter family member 74D-like in the midgut, vesicle transport through interaction with t-SNAREs homolog 1B-like in the ovaries, enteropeptidase-like in the salivary glands, acetylcholine receptor subunit alpha-like in the synganglion, and cuticle protein 19.8-like in the trachea (Supplementary Table S7). Similarly, no clear tissue association was found when cluster-level gene expression profiles were compared to the reference atlas (Fig. 10B; Fig. S3). Overall, we do not detect clear-tissue specific signatures in the ISE6 cell line. This supports the conclusion that this cell line is transcriptionally diverse but not committed to any defined tissue lineage.

**Figure 10.**
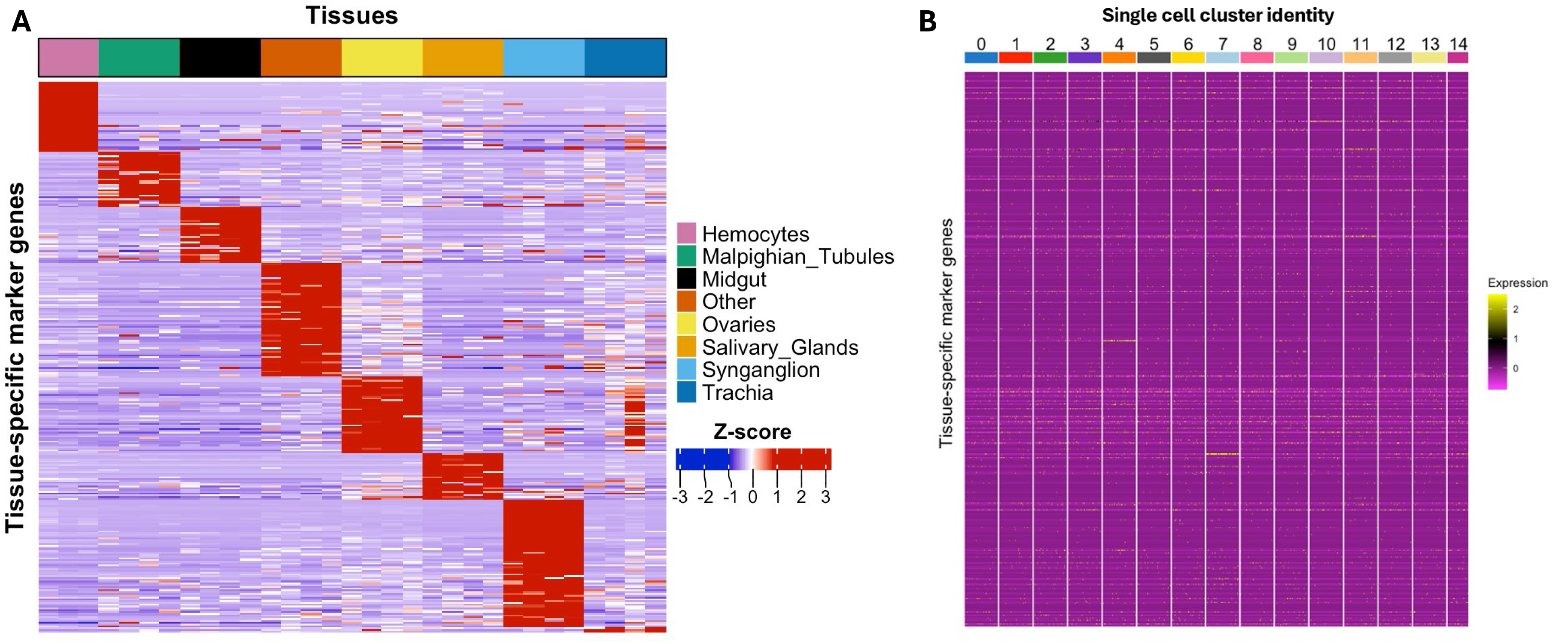
Comparison of cell and tissue specific transcriptional signatures in bulk tissue RNA sequencing and single-cell RNA sequencing of ISE6 cells. (A) Heatmap showing Z-score–normalized expression of marker genes associated with multiple tick tissues and carcasses (called “others”), including the top expressed marker genes for each tissue category, across ISE6 cells. (B) Heatmap illustrating the normalized expression patterns of tissue-associated marker genes across the fifteen cell clusters identified by single-cell RNA sequencing analysis, indicating the absence of clearly defined tissue- or cell-type–specific transcriptional signatures within the ISE6 cell population.

## DISCUSSION

In this study, we demonstrate that EME infection of ISE6 cells induces a progressive, time-dependent transcriptional response characterized by early stress responses followed by downregulation of metabolic and proliferative pathways. Single-cell analysis further reveals that this response occurs across a morphologically and transcriptionally heterogeneous cell population. Together, these findings clarify host cellular responses during intracellular ehrlichial infection and provide new insight into the biological behavior of the ISE6 model system. Although we observed differences in ISE6 cell morphology, EME did not preferentially infect any specific cell population. Morulae were observed across morphologically distinct cells, and bacterial replication correlated with progressive decline in host cell viability. This pattern is consistent with previous observations of EME in multiple tick tissues during feeding (Lynn et al., 2019; Aspinwall et al., 2025). Rather than targeting a specific cell type, EME could enter and replicate in multiple cell types, suggesting non-specificity in its cellular niche. The decline in viability observed during infection aligns with prior ultrastructural studies of ehrlichial pathogens in both tick cells and tissues and mammalian cells resulting in cytoplasmic distortion of cells and pronounced distortion and lysis of tissues (Popov et al., 2007; Dedonder et al., 2012; Munderloh et al., 2014; Popov et al., 1995). Host-cell viability outcomes can differ across tick-borne pathogens and experimental systems, including variation by cell line, pathogen, and inoculation conditions (Zivkovic et al., 2009; Kuriakose et al., 2011; Ruzek et al., 2008; Weisheit et al., 2015; Mansfield et al., 2017), supporting the idea that the late-stage decline we observe could reflects a combination of increased EME burden and cell-line-specific susceptibility.

At early point, EME infection was characterized predominantly by upregulation of genes associated with unfolded protein response, mitochondrial organization, redox balance, and TOR signaling. This early response is consistent with pathogen-associated stress responses previously reported for other tick-pathogen, including *A. phagocytophilum* in *Ixodes* cell lines and tick-borne encephalitis virus infection (Alberdi et al., 2016; Grabowski et al., 2016). The induction of the ER chaperone BiP and IRE1-associated pathways is consistent with conserved ER stress responses (Schröder et al., 2005).

This is consistent with observations made during *Anaplasma phagocytophilum (A. phagocytophilum)* and *Borrelia burgdorferi (B. burgdorferi)* infections, where the unfolded protein response modulates immune signaling in ticks by activating the immune-mediated deficiency (IMD) pathway of *I. scapularis* to restrict *A. phagocytophilum* and *B. burgdorferi* (Sidak-Loftis et al., 2022). The increased peroxiredoxin expression is consistent with adaptation to a highly oxidative environment such as high iron production which generates a high amount of reactive oxygen species (ROS) during blood feeding (Citelli et al., 2007; Galay et al., 2015). Iron is also essential for the survival and proliferation of ehrlichial pathogens (Luo and McBride, 2012; Moumène et al., 2018). Recent studies demonstrated that *E. chaffeensis* hijack host cell iron stores by inducing ferritinophagy, thereby increasing the labile iron pool required for bacterial growth (Yan et al., 2018; 2021). The interplay between iron acquisition, oxidative stress, and antioxidant defenses appears to be a conserved strategy among ehrlichial pathogens for successful colonization and transmission (Barnewall & Rikihisa, 1994; Barnewall et al., 1999; Yan et al., 2021). To counteract infection-induced oxidative stress, ticks rely on antioxidant enzymes such as peroxiredoxins, catalases, and the selenocysteine insertion sequence binding protein 2 (SBP2), which is essential for selenoprotein-mediated redox regulation. Beyond protecting host tissues from ROS, these antioxidant systems have been shown to facilitate pathogen colonization and transmission. Enhanced antioxidant activity in *Amblyomma maculatum* promotes vertical transmission of *Rickettsia parkeri* (Budachetri et al., 2017), and catalase has been demonstrated to be required for efficient arbovirus colonization in *Aedes aegypti* (Oliveira et al., 2017). Similar redox-regulatory mechanisms have also been implicated in tick responses to *Anaplasma* and *Rickettsia* infections (Alberdi et al., 2019; Dahmani et al., 2020). The upregulation of peroxiredoxin observed here likely reflects a response to EME-induced ROS generation and may contribute to creating a redox-balanced intracellular environment that supports pathogen survival.

Concurrently, downregulation of mismatch repair components suggests EME-driven modulation of DNA maintenance pathways during early infection. While functional consequences of reduced MSH2, MSH6, and PCNA expression are not established in arthropod cells, analogous systems indicate that altered mismatch repair capacity can influence stress tolerance (Grazielle-Silva et al., 2015; 2018), and evolutionary analyses emphasize mismatch repair as a major determinant of genome stability and mutational dynamics (Muthye and Lavrov, 2021). This is particularly notable because survival of *B. burgdorferi* in ticks under oxidative/nitrosative stress depends on intact DNA repair pathways (Bourret et al., 2016), and comparative work on *Midichloria mitochondrii* during infection of *I. ricinus* and *I. holocyclus*, indicates that stress can drive distinct DNA repair strategies (Leclerc et al., 2024). Whether mismatch repair downregulation during EME infection reflects pathogen-driven manipulation or a secondary consequence of cellular stress remains to be determined.

As infection progressed, transcriptional activation transitioned into widespread repression. By day 4, genes involved in kinase signaling, lipid metabolism, mitochondrial regulation, DNA replication, and cell cycle progression were broadly downregulated. The coordinated suppression of replication machinery, checkpoint regulators, and cytoskeletal components indicates marked attenuation of proliferative capacity. In contrast to what was previously reported in vivo, where genes involved in metabolism, immune defense, histone modification, and oxidative stress responses were markedly upregulated in *A. hebraeum* midguts during *E. ruminantium* infection (Omondi et al., 2023). Similar metabolic reprogramming has been described for *A. phagocytophilum* in ISE6 cells, where increased mitochondrial Phosphoenolpyruvate carboxykinase (PEPCK) activity enhances phosphoenolpyruvate (PEP) production while cytosolic PEPCK is downregulated, promoting intracellular PEP accumulation without overt loss of host cell viability (Villar et al., 2015; Cabezas-Cruz et al., 2017). This metabolic rewiring appears to support persistent infection while maintaining host cellular function. These findings raise the possibility that intracellular bacteria may differentially alter host metabolism even in the same cell, with some pathogens maintaining metabolic homeostasis to sustain persistence, whereas others induce longer-term transcriptional repression that limits host proliferative capacity.

The mitochondrial-related signatures observed early in EME infection are consistent with studies showing that obligate intracellular pathogens modulate host mitochondrial dynamics and apoptosis (Xiong et al., 2008; Liu et al., 2012). In our dataset, early upregulation of mitochondrial fission 1 protein (FIS1) and prohibitin-2 suggests remodeling of mitochondrial quality control pathways. FIS1 regulates mitochondrial fission, a process linked to mitophagy and stress-induced apoptosis (Twig and Shirihai, 2011), while prohibitin-2 contributes to mitochondrial integrity and apoptotic regulation. Ehrlichial pathogens have been shown to directly manipulate mitochondrial function. In mammalian cells, *E. chaffeensis* recruit mitochondria and delays apoptosis through mitochondrial-targeted effectors, which inhibit Bax-mediated apoptosis and modulates ROS, thus facilitating bacterial replication and infection (Liu et al., 2012). Mitochondrial remodeling is also recognized as a common feature of host–microbe interactions in arthropods, including viral induction of mitophagy seen with dengue virus infection in mosquito (Santana-Román et al., 2021) and symbiont-driven mitochondrial fragmentation in the case of *Midichloria mitochondrii* infection in the *I. ricinus* ticks (Uzum et al., 2023).

The ISE6 cell line has served as a foundational tool for studying tick–pathogen interactions for more than three decades (Munderloh et al., 1994; Kurtti et al., 1996; Singh et al., 2024). However, its cellular identity and transcriptional organization have remained incompletely defined. In this study, single-cell transcriptomic analysis reveals that ISE6 cells comprise a morphological and transcriptionally heterogeneous population that lacks clear tissue-specific differentiation. Furthermore, EME infection induces a progressive, time-dependent shift from early stress adaptation to late-stage transcriptional and proliferative suppression. Together, these findings support the biological interpretation of ISE6 cells as a model system and clarify host cellular responses during intracellular ehrlichial infection.

Both ultrastructural and single-cell analyses demonstrate substantial heterogeneity within ISE6 cultures. This variability is consistent with the embryonic origin of the cell line, which was derived from developing eggs containing multiple differentiating cell populations (Munderloh et al., 1994; Hinne et al., 2025). Embryogenesis in ticks involves dynamic and asynchronous differentiation of multiple cell populations, and comparable developmental studies in other tick species similarly highlight the establishment of diverse cell types during early development (Campos et al., 2006; Campos et al., 2007; Santos et al., 2013; Friesen et al., 2016). Prior work has suggested that ISE6 cells can exhibit hemocyte-like morphology and mixed epithelial or neuronal features (Villar et al., 2015; Oliver et al., 2015; Alberdi et al., 2016; Cabezas-Cruz et al., 2017). In contrast, our single-cell data does not support similarity to defined adult tissues. Even when mapped against a comprehensive tissue reference atlas, clusters failed to demonstrate enrichment for midgut, salivary gland, synganglion, or hemocyte-specific signatures (Rolandelli et al., 2024). These findings indicate that transcriptional variation within ISE6 reflects functional states rather than tissue-type commitment. This distinction is important, as prolonged in vitro culture and selective pressures can drive divergence from native tissue identities or specific selection of certain cell types, a phenomenon well documented in arthropod and insect cell lines (Krzywinska et al., 2022).

Although ISE6 clusters did not map specific tissues, several cluster markers align with potential functions previously implicated in tick physiology. Clusters containing transcripts expressing neuropeptide receptor and ion transport are consistent with neuroendocrine or excitability-associated regulation: pyrokinin signaling has documented roles in gut motility and ion/water balance in *Aedes aegypti* (Lajevardi and Paluzzi, 2020), while sodium/calcium exchangers contribute to calcium homeostasis in excitable cells in *Drosophila* (Oberwinkler and Stavenga, 2000). Similarly, extracellular matrix-associated genes such as hemicentin family members are linked to tissue contact and migration in *Caenorhabditis elegans* (Gianakas et al., 2023). The presence of function specific transcripts without tissue type overlap suggests that ISE6 cells retain functionally relevant gene repertoire shared across tick tissues, even if it does not recapitulate discrete in vivo differentiation states. Supporting this, many transcripts present in ISE6 overlap with genes reported in tick midgut and salivary gland transcriptomic or proteomic studies (Schwarz et al., 2014; Lu et al., 2023), reinforcing the continued utility of ISE6 as a tractable experimental system and as a platform for isolating and propagating tick-borne pathogens (Moniuszko et al., 2014).

In conclusion, we demonstrate that EME infection in ISE6 cells induces a time-dependent transcriptional response characterized by early activation of stress and metabolic adaptation pathways, followed by progressive restriction of key signaling and proliferative programs as bacterial replication increases. Despite this dynamic response, EME did not preferentially infect a specific cell population. Rather, the ISE6 cell line comprises a morphologically and transcriptionally heterogeneous mixture of cells, and these differences were not associated with selective susceptibility to infection. Further mechanistic studies are needed to define the functional overlap between the different cell populations present in the ISE6 cell line and the cell types found in distinct tick tissues. This information will strengthen the utility of tick cell lines, such as ISE6, for investigating tick biology and tick–pathogen interactions.

## ACKNOWLEDGEMENTS

The authors want to offer special thanks to the NCI sequencing facility at NIH for performing the sequencing of the samples. We also acknowledge Drs Lucas Tirloni, Larissa A. Martins and Diane Cockrell for the initial discussions.

## Author Contributions

Study design and conceptualization: TBS, JA, AA; Experiments execution and methodology: TBS, JA, FH, VN; Data analysis: TBS, AA, JA, CM, AM, MCWH, VN, JL; Manuscript preparation and revisions: TBS, AA, JA, CM, OO, RR, FH, VN, JL; Funding acquisition: TBS, CS; Supervision: TBS.

## Data and materials availability

The sequencing data generated in this study have been deposited in the NCBI Sequence Read Archive (SRA) under BioProject accession PRJNA1416822.

## SUPPLEMENTAL MATERIALS

**Supplemental Text.** This section provides additional methodological details and extended results, including single-cell RNA sequencing data processing, differential gene expression analyses, Gene Ontology enrichment results, and tissue-specific gene identification. The primary essential information supporting the main findings is presented in the main text.

**Supplemental Figure S1.** Gating strategy for flow cytometry analysis. (A) Identification of whole cells using SSC-A versus FSC-A. (B) Identification of singlets using FSC-A versus FSC-H. (C) Identification of viable cells using Live/Dead DAPI nuclear stain. (D) Live cells were gated for EME infection using fluorescently labelled α-*Ehrlichia* OMP-19 specific antibodies. FSC-A, Forward Scatter Area; FSC-H, Forward Scatter Height; SSC-A, Side Scatter Area.

**Supplemental Figure S2.** Single-cell quality metrics across experimental groups, showing (A) the number of genes detected per cell (nFeature_RNA) and (B) the percentage of mitochondrial genes (% mitochondrial) in EME_d2, EME_d4, UI_d2, and UI_d4 ISE6 cells.

**Supplemental Figure S3.** (A) Principal component analysis (PCA) of the sc-RNA seq gene expression datasets. The points of the scatter plots visualize the samples based on infection status and time points. The colors represent samples from four different datasets, respectively. (B) Proportion of cells from uninfected and EME-infected ISE6 cells at day 2 and day 4 p.i. within each unsupervised cluster.

**Supplemental Table S1**. Putative marker genes identified for each of the 15 transcriptional clusters in ISE6 cells, including differential expression analysis between uninfected and EME-infected conditions at day 2 and day 4 p.i.

**Supplementary Table S2**. Differentially expressed genes (DEGs) identified across pairwise comparisons between uninfected and EME-infected ISE6 cells at day 2 and day 4 p.i.

**Supplementary Table S3**. Gene Ontology (GO) enrichment analysis of differentially expressed genes in EME-infected ISE6 cells at day 2 p.i. compared to uninfected controls.

**Supplementary Table S4**. Gene Ontology (GO) enrichment analysis comparing EME-infected ISE6 cells on day 4 and day 2 p.i., highlighting pathways associated with infection progression.

**Supplementary Table S5**. Gene Ontology (GO) enrichment analysis of differentially expressed genes in EME-infected ISE6 cells at day 4 p.i. compared to uninfected controls.

**Supplementary Table S6**. Transcription factor expression analysis across ISE6 single-cell clusters, including composite transcriptional scores and differential regulation between uninfected and EME-infected conditions.

**Supplementary Table S7**. Tissue-specific marker genes identified from bulk RNA sequencing of *Ixodes scapularis* tissues and published hemocyte datasets, used to construct the reference transcriptional atlas and assess tissue signature enrichment in ISE6 single-cell clusters.

